# Intracellular label-free detection of mesenchymal stem cell metabolism within a perivascular niche-on-a-chip

**DOI:** 10.1101/2020.10.03.322297

**Authors:** Simone Perottoni, Nuno G. B. Neto, Cesare Di Nitto, Manuela Teresa Raimondi, Michael G. Monaghan

## Abstract

The stem cell niche at the perivascular space in human tissue plays a pivotal role in dictating the overall fate of stem cells within it. Mesenchymal stem cells (MSCs), in particular, experience influential microenvironmental conditions, which induce specific metabolic profiles that affect processes such as cell differentiation and dysregulation of the immunomodulatory funtion. Reports focusing specifically on the metabolic status of MSCs under the effect of pathophysiological stimuli - in terms of flow velocities, shear stresses or oxygen tension – do not model heterogenous gradients, highlighting the need of more advanced models reproducing the metabolic niche. Organ-on-a-chip technology offers the most advanced tools for stem cell niche modelling thus allowing for controlled dynamic culture conditions while profiling tunable oxygen tension gradients. However, current systems for live cell detection of metabolic activity inside microfluidic devices require the integration of microsensors that allow for extracellular measurments only, giving innacurate and indirect information about the metabolic state of cells. Here, we present a metabolic toolbox coupling a miniatuirzed *in vitro* system for human-MSCs dynamic culture, that mimics microenvironmental conditions of the perivascular niche, with high-resolution imaging of intracellular metabolism. Using Fluorescence Lifetime Imaging Microscopy (FLIM) we monitor the spatial metabolic machinery and correlate it with experimentally validated intracellular oxygen concentration after designing the oxygen tension decay along the fluidic chamber by *in silico* models prediction. Our platform allows for the subjection of a metabolic profile to MSCs, mimicking the physiological niche in space and time, and its real-time monitoring representing a functional tool for modelling perivascular niches, relevant diseases and metabolic-related uptake of pharmaceuticals.

## Introduction

The perivascular spaces of many organs contain adult progenitor cell populations in a microenvironment called the perivascular stem cell niche, that presents specific conditions to maintain multi-lineage potential and self-renewal capacity.^1,2^ This is a complex and dynamic milieu, where stem cells respond to specific biochemical and mechanical cues. The perivascular niche is an open system, inhabited by heterogeneous cellular populations while physicochemical factors such as oxygen concentration, nutrient availability, signalling factors as well as pH, shear stress or temperature, contribute to its homeostasis.^3^ Dysregulation of the niche’s microenvironment, associated with hypoxia, inflammation or metabolic reprogramming, has recently been shown to elicit niche-related pathologies.^4,5^ The influence of cellular metabolism on the overall fate of cells is eliciting a specific interest. It has long been established that cellular metabolism is intrinsically linked to cell behaviour and phenotypic status; with one of the earliest historical reports in 1924 detailing the Warburg Effect exhibited by cancer cells when instates of aerobic and anaerobic metabolism.^6^ Only recently has the appreciation of cellular metabolism begun to gather attention in the realms of stem cells; with demonstrations of a pivotal role of the metabolic state in accompanying cell fate towards proliferation, quiescence or differentiation. Shifts in the balance between a glycolytic metabolic profile and the mitochondrial oxidative phosphorylation (OxPhos) define the contribution of adult progenitor and stem cells to the renewal and homeostasis of their native tissue.^7–10^ Among the different cell populations involved in niche homeostasis, Mesenchymal stem cells (MSCs), which are multipotent cells present in all vascularized tissues ^11^; are emerging as key players in the modulation of the overall niche response and organization.^12,13^

Moreover, due to their immunomodulatory properties, human-MSCs (h-MSCs) have been extensively applied as therapeutic agents.^14,15^ However, *in vitro* manipulation is shown to alter cell metabolism influencing h-MSC’s functional properties during *in vitro* expansion.^16^ Several reports have examined the effect of flow, shear stress and pressure on stem cell differentiation and mineralisation, but reports specifically focusing on the metabolic status of cells in such conditions are rather lacking.^17–20^ Despite efforts in improving standard 2D culture protocols ^21–23^, advanced tools for controlled and physiologically relevant *in vitro* cultures of h-MSCs are still necessary for enhancing knowledge on their metabolic behaviour both *in vivo* and *ex vivo*. Advanced *in vitro* models of the stem cell niche offer the possibility of studying directly and manipulating human stem cell populations, within biomimetic microenvironments, with the potential of providing valid alternatives to animal models in studying tissue renewal and pathogenesis.^24^ Such niche-on-a-chip (NOC) systems include fluidic devices that recapitulate relevant feature of the *in vivo* niche microsystem within fluidic channels for dynamic cell culture.^25–28^ These systems can also facilitate defined mechanical stimulation (in terms of shear forces) and spatiotemporal control of biomolecular transport by recapitulating flow velocities at interstitial levels.

However, these systems do possess important limitations in their ability to reproduce the mechanical and mass transport field variables present in a native perivascular niche. 1) Microfluidic bioreactors can be utilised to generate a controlled cellular metabolism capitulating that present in the perivascular niche, by the engineering of a defined oxygen tension gradient ^29–31^ that mimics the spatial oxygenation at the interface between blood vessels and the stem cell niche (figure 1).^32,33^ Standard 2D cell culture methods are limited in that they typically impose uniform static normoxic or hypoxic oxygen environments. 2) Cellular metabolism *in vitro* is usually assessed using end-point staining assays which provide intracellular quantification but are an end-point assessment that requires culture sacrifice. 3) Microsensor integration within organ-on-chip platforms can enable continuous metabolic monitoring and recording by accessing the specific molecules involved in cellular metabolism. However, such sensors often operate via enzymatic or biochemical mechanisms which can influence the cellular microenvironment and respiration. The dimensions of currently available microsensors confines their use to extracellular readings, meaning that they are limited to indirect assessment of global cellular metabolism.^34^

**Figure 1.**
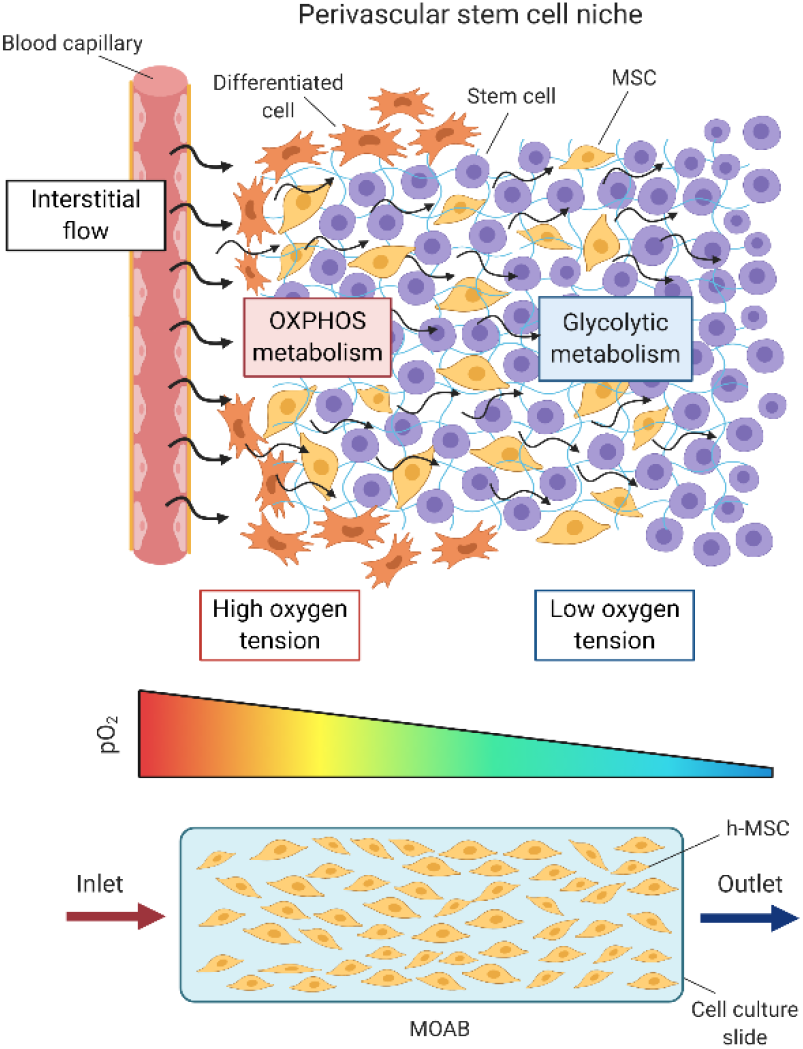
Mesenchymal stem cells (MSCs) reside within perivascular spaces called perivascular niches, which exert regulatory control on the stem cells by interstitial flow levels and defined oxygen tension gradients leading to a specific spatial metabolic profile.^8^

Advanced imaging technologies are becoming valuable tools for live label-free assessment of intracellular behaviour, and can be applied for metabolic profiling at a single-cell level.^7^ Among the available fluorescence microscopy methods, Fluorescence Lifetime Imaging Microscopy (FLIM) has emerged as a key technique capture interaction of specific biomolecules in living cells. It provides high photon efficiency, lifetime accuracy, real-time measurements, high spatial resolution, and does not require staining or labelling of fluorophores. Widespread FLIM applications consist in mapping intracellular temperature, viscosity, pH as well as measuring ions, glucose or oxygen concentration. FLIM can capture the signals emitted by endogenous fluorophores such as NAD(P)H or FAD, providing a readout of the metabolic state of the samples under investigation.^35,36^

Here, we present a metabolic toolbox that couples a micro-physiological system (MPS) for h-MSCs dynamic culture capable of simultaneous imaging of intracellular metabolism at a high resolution. Predictive modelling was employed to engineer a declining gradient in oxygen tension and validated using a previously calibrated non-invasive imaging technology applicable to this miniaturised optically accessible bioreactor (MOAB). Using FLIM we were able to monitor the preferential modes of metabolic machinery employed in addition to visualising the correlating oxygen concentration available at these points which were in agreement with our *in silico* model. These results present a new method and technology to preferentially pattern an in silico-modelled metabolic gradient which will be of enormous benefit to the community in modelling perivascular niches, the relevant diseases, and also the bioavailability and metabolism-related uptake of pharmaceuticals.

## Results and discussion

### The niche-on-a-chip (NOC) for stem cell metabolism measurement

#### The microbioreactor for h-MSCs culture

The culture platform employed in this study is a miniaturized optically accessible bioreactor (MOAB), recently reported for culturing h-MSCs inside a dynamic MPS ^37^ (figure 2A-B).

**Figure 1.**
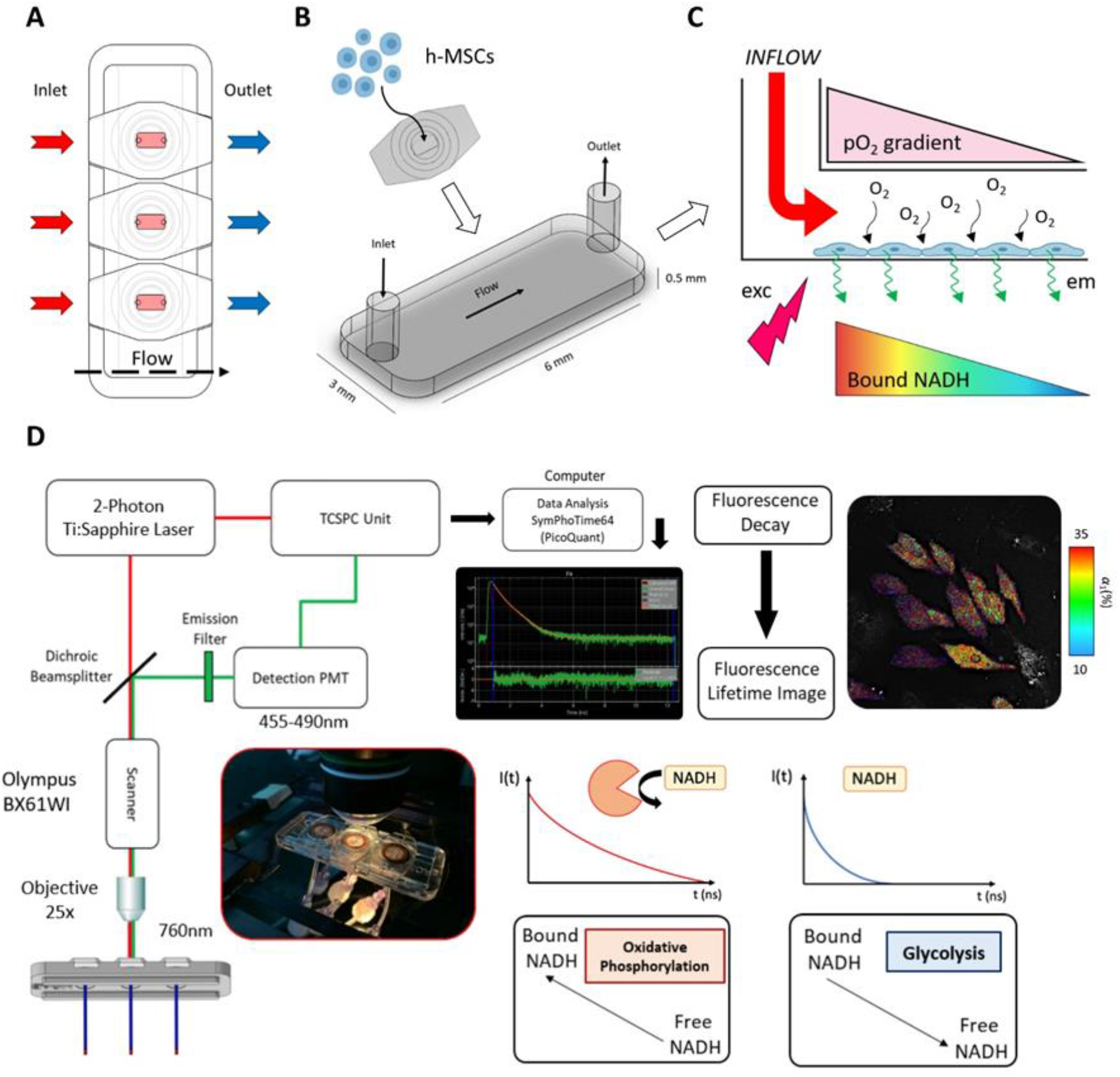
Microfluidic bioreactor and experimental setup. (A) The Miniaturized Optically Accessible Bioreactor (MOAB) device, composed of a Polycarbonate (PC) body with three independent rectangular base flow chambers for cell culture and real-time high-resolution imaging. (B) “Plug and play” seeding of human-MSCs (h-MSCs) inside the fluidic chamber for interstitial - like perfusion. (C) By using MOAB a defined oxygen tension gradiant can be imposed by controlling the flow perfusion and cell density while simultaneous real-time high-resolution measurement of the metabolic activity by the detection of the bound NAD(P)H fluorescence emission can be performed. {D) Non-invasive measurement of cellular metabolism of h-MSCs cultured inside MOAB, using a two-photon (2P) microscope fitted with Fluorescence Lifetime Imaging Microscopy (FLIM) detectors. The result is the NAD(P)H fluorescence decay and a colour coded picture indicating pools of free and enzyme-bound NAD(P)H quantified by their respective fluorescence lifetimes fractions. The free NAD(P)H corresponds to the short τ_1_ lifetime with corresponding fraction α_1_, representative of a glycolytic profile, while the protein bound NAD(P)H is related with the long τ_2_, and α_1_, fraction, indicating oxidative phosphorylation (OxPhos).

Perfused *in vitro* cultures of h-MSCs have demonstrated the influence of flow-associated shear stress on MSCs commitment to the mesengenic lineages.^38,39^ Here, precise control enables interstitial flow velocities and low level shear forces as well as generate a transitional oxygen concentration profile in space and time with the aim of mimicking an interface representative of the one that h-MSCs experience near blood capillaries and within the stem cell niche.

The walls of the MOAB microchambers are fabricated from oxygen-impermeable polycarbonate (PC, with an oxygen diffusion coefficient of 8.0 × 10^−8^cm^2^ s^−1^) ^31^ thus enabling a stable and linear gradient of the oxygen tension. Indeed, the only source for free oxygen to reach inside the microchamber is the oxygenated culture medium flowing from the inlet. The combination of constant flow of fresh medium with the consumption of oxygen by the cells modulates an oxygen tension in the microenvironment, leading to the establishment of a spatial oxygen gradient (figure 2C).

#### Intracellular label-free investigation of h-MSCs metabolism

Transparent PDMS microfluidic chips permit non-invasive optical monitoring of the MOAB microchamber for the duration of the experimentation. However, in order to perform high-resolution imaging, immersion objectives are necessary and the refractive index (n) of the viewing surface needs to be as close as possible to that of the objectives. PDMS (with n=1.43) generates optical aberrations on the excitation laser pathway; therefore, the overall optical quality is generally poor. Moreover, PDMS membrane thickness cannot be reduced substantially without interfering with the performance of the device.^40^ We integrated a rectangular glass slide (having n = 1.53 and a thickness of 0.15 mm) inside each fluidic chamber that simultaneously serves as a surface for cellular adhesion to enable high-resolution microscopy. This optical setup facilitates real-time visualisation of cellular metabolism by means of a two-photon (2P) microscope fitted with FLIM detectors (Fig. 2C).

Harvesting the autofluorescence properties of NAD(P)H provides an insight to the metabolic state of cells.^41–43^ This is based on the presence of mixed pools of free and enzyme-bound NAD(P)H that can be individually discernible by their respective fluorescence lifetimes. Here, we focused on the use of 2P-NAD(P)H FLIM. NAD(P)H is characterised by a multi-exponential fluorescence decay, which is fitted using a double-exponential curve with an average fluorescence lifetime (τavg) obtained from the two different states fluorescence lifetimes and corresponding fractions (τ1-2,α1-2).^44–47^ A representative NAD(P)H fluorescence decay, its fitting and residuals are shown in Fig. 2D. The free NAD(P)H corresponds to the short τ1 lifetime (~0.4 ns) with corresponding fraction α1 while the protein-bound NAD(P)H is related with the long τ2 (>1.5 ns) and α2 fraction ^47–49^. NAD(P)H is a co-factor of important steps of the metabolic pathways, including glycolysis, OxPhos and pentose phosphate pathway ^50^.

#### Optimization of the MPS culture conditions

The spatial and temporal distribution of the oxygen tension across a microfluidic chamber depends on the uniformity of the flow field and on the oxygen consumption rate of cells exposed to this flow. A computational approach was employed to establish a defined perfusion regime at the inlet to create a specific oxygen gradient along the principal flow direction, while ensuring physiological physical stimulation to cells.

#### Prediction of interstitial flow velocities and shear stress

CFD analysis (COMSOL Multiphysics^®^) validated in previous work ^27^ predicted the velocity distribution profile and the levels of shear stress acting in the MOAB microchamber. Interstitial flow velocities in body tissues generate from convective transvascular flows and are reported to range within 0.1-5 μm s^−1^, in physiological conditions, reaching up to 10 μm s^−1^ during inflammation.^51^ Four flow rates (0.5, 2, 3.5 and 5 μL min^−1^) were simulated, yielding average velocity values in the range 0.5-6 μm s^−1^ (figure 3A).

**Figure 3.**
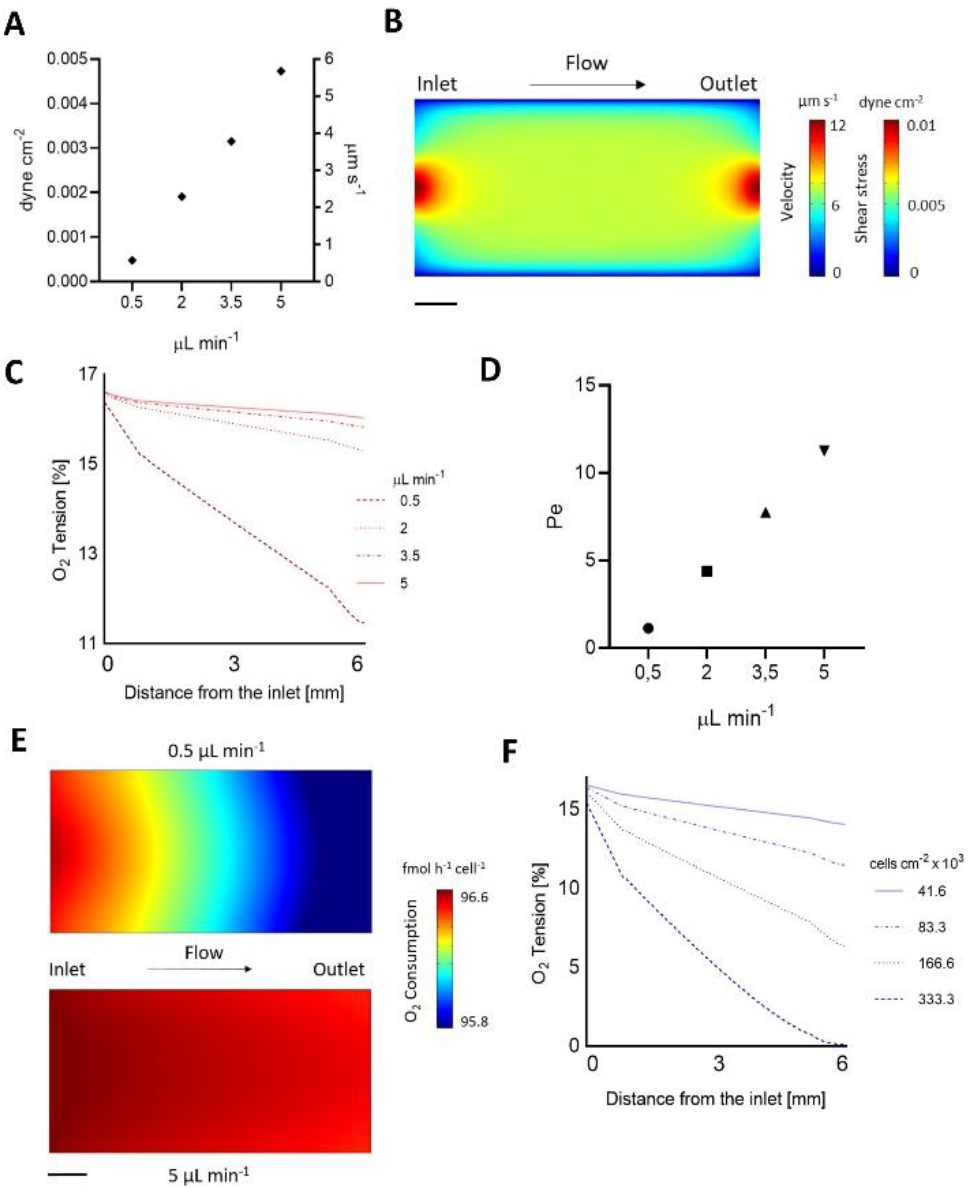
Velocity profile, shear stress and oxygen tension inside the perivascular NOC. (A) Predicted average flow velocities and wall shear stress as function of the input flowrate. (B) Colour map of the flow velocity and wall shear stress distribution in proximity of the cell culture surface area at an inlet flowrate of 5 μL min^−1^. Colour map in top view. (Scale bar: 1mm.) (C) Prediction of the oxygen tension depletion at a cell density of 8.3 × 10^4^ cells cm^−2^ (corresponding to 96 hours of culture) as function of the distance from the inlet and the input flowrate. (D) Computed P*e* number as a function of flowrate considering an oxygen particle at the middle height of the chamber (characteristic length L: 250 μm) in correspondence of the max velocity fillet on a section at 3 mm distance from the inlet. (E) Colour maps of the single cell oxygen uptake rate spatial gradient at 0.5 μL min^−1^ (top) and 5 μL min^−1^ (bottom) at a cell density of 8.3 × 10^4^ cells cm-2 (corresponding to 96 hours of culture). Colour maps in top view. (Scale bar: 1 mm.) (F) Prediction of the oxygen tension depletion at a flowrate of 5 μL min^−1^ as function of the distance from the inlet and the cell density (i.e. the day of culture).

The estimated wall shear stress, resulting from the flow rates applied (0.5, 2, 3.5 and 5 μL min^−1^), ranged from an average value of 0.0005 dyne cm^−2^, at the lowest flowrate (0.5 μL min^−1^), to 0.005 dyne cm^−2^ at the highest (5 μL min^−1^) (figure 3A). The highest shear stress predicted to act on cells reached 0.001 dyne cm^−2^ which is orders of magnitudes lower than the values reported for shear stress-induced MSCs differentiation.^20,38,39^ The velocity profile together with shear stress were uniformly distributed at the bottom of the culture chamber with 75% of the surface simulating values within a 10% variation (Fig. 3B). This simulation predicts equal mechanical stimulation (in terms of shear stress) to the cells.

#### Flow-dependent oxygen concentration profile

Next, a mass transport analysis was implemented (COMSOL Multiphysics^®^) to predict the oxygen tension profile with consideration of h-MSCs oxygen consumption across the culture chamber. The cellular consumption rate was modelled as an oxygen sink by defining a steady state flux, dependent on cell density, along the cell adhesion surface of the MOAB chamber while assuming a uniform distribution of cells. Single cell consumption rate was assumed to depend on both the cell type and on local oxygen availability, following the Michaelis-Menten (MM) equation.^52^ As a boundary condition we imposed an oxygen partial pressure of 16.7% at the inlet of the MOAB chamber, based on experimental measurements.^53^

From this simulation the oxygen concentration profile gradually decreased across the chamber given a fixed cell number, and its gradient rapidly decreased with flow velocity (Fig. 3C). At high flow velocity, the contribution of advection prevailed on diffusion within the medium; and in this case it was represented by a 90% increase of the Peclet number (Pe) at the highest flow rate (Fig. 3D). Moreover, at a fixed reference cell density of 6 × 10^4^ cells cm^−2^, the oxygen tension exponentially decreased with the flowrate due to the h-MSCs oxygen uptake. Simulating cellular O_2_ consumption decays at the two opposite flowrates (i.e. 0.5 and 5 μL min^−1^) highlights diverse oxygenated microenvironments as function of the distance from the inlet (Fig. 3E).

Besides the flow velocity, the other element influencing the overall oxygen tension profile is the cell density. We investigated four cell density values (41.6, 83.3, 166.6 and 333.3 cells cm^−2^ × 10^3^), representing the cells doubling during the days of culture, at one fixed flowrate of 0.5 μL min^−1,^ which is the lowest in the MOAB working range. At every cell doubling, the oxygen tension gradient got steeper generating always more hypoxic conditions at the chamber outlet. After four doublings, the oxygen tension at the outlet reached 0%, representing the upper limit working condition of our system in time. This computational approach informed the preparation of the optimal experimental working conditions. The numerical results demonstrated the feasibility to simulate *in vitro* the varying intensities at which fluids extravasate into the stem cell interstitium, due to changes in blood vessel permeability.

These simulations allowed us to predict oxygen concentration by delineating the effect of varying flowrates on oxygen diffusion and the impact of cell density on the oxygen availability thus, on the overall normoxic-hypoxic gradient within this MPS.

Oxygenation levels within different body tissues and organs vary greatly. For example, the oxygen tension in lungs and liver ranges from 13% to 10% while in bone marrow or brain it reaches values from 7% down to 4%.^29,32,33^ By tuning the input flow parameters in the MOAB, we were able to recapitulate specific physiological oxygenation levels and oxygen concentration decay. This represents a functional tool that could be useful in modelling the perivascular niche present in various organs, besides the bone marrow niche modelled here, which is the one primarily investigated to elucidate its role in MSCs differentiation.^11,54,55^

#### Quantitative intracellular oxygen measurement

To assess and validate the predicted cell oxygen uptake gradient being formed inside of the MOAB, we sought to experimentally quantify intracellular oxygen. To ensure the most physiologically relevant measurements, we performed fluorescence intensity based measurements inside the MOAB using an intracellular fluorescence probe ^56^. Specifically, we used a SI-0.2^+^ fluorescence probe composed of fluorine building blocks which act as an energy donor and an antenna for platinum(II) meso-bis(pentafluorophenyl)bis(4-bromophenyl)porphyrin (PtTFPPBr2) ^57^. This conjugated polymer exhibits favourable properties to be used a multiphoton system due to its high intensity fluorescence emission, cell permeability and Förster resonance energy transfer (FRET) properties which is impacted in varying oxygen concentrations. At higher oxygen concentrations, this probe is quenched and lower amounts of energy is transferred from the backbone to metalloporphyrine complex resulting in a decrease of emission intensity at 640-680 nm.

After h-MSCs incubation with the probe for 16 hours, it was possible to establish a calibration curve (supplemental figure 1) by detecting the fluorescence emission intensity in cases of uptake activity in normoxic conditions and uptake when a compromised cellular oxygen consumption rate was engineered by suppression of cells respiratory activity.

h-MSCs were cultured inside the MOAB for 96 hours at a flow rate of 0.5 μL min^−1^. Afterwards, they were stained using SI-0.2^+^ and imaged directly in the perfused chamber with a multiphoton microscope while maintaining the same flow rate. Three areas of the MOAB were analysed: inlet, middle and outlet (figure 4A). Two images were acquired at the same time, one for NAD(P)H emission (as a reference signal) and another for the probe emission wavelength. Afterwards, the emission intensity in the cell area was calculated and a ratio of both acquisitions was performed. A statistically significant increase in the ratio 630/460 from the inlet area of 1.09 ± 0.04, to 1.44 ± 0.10 in the middle, and finally 1.68 ± 0.17 in the outlet area (figure 4B) was detected. As described previously, an increase of the 630/460 ratio is correlated with decreasing oxygen levels due to decrease of the efficiency of energy transfer from the fluorine backbone to the metalloporphyrin ^57^. By applying our semi-calibration curve (supplemental), it was possible to correlate the intensity measurements of the 630/460 ratio to partial oxygen values and obtain an estimation of the intracellular oxygen levels. Here, we calculated that the inlet oxygen levels are on average 16.6 ± 0.4 %, 13.1 ± 1.0 % in the middle and 10.7 ± 1.7 % while maintaining their statistical difference (figure 4C). Therefore, we were able to observe a direct effect on the intracellular oxygen levels promoted by the flow rate and cell culture environment. This result agrees with the calculations of the *in silico* model (figure 3). Here, a similar decreasing trend in oxygen distribution from the inlet (≈16.5%) to the outlet (≈ 11.5%) region is calculated in our model (figure 3C).

**Fig. 4.**
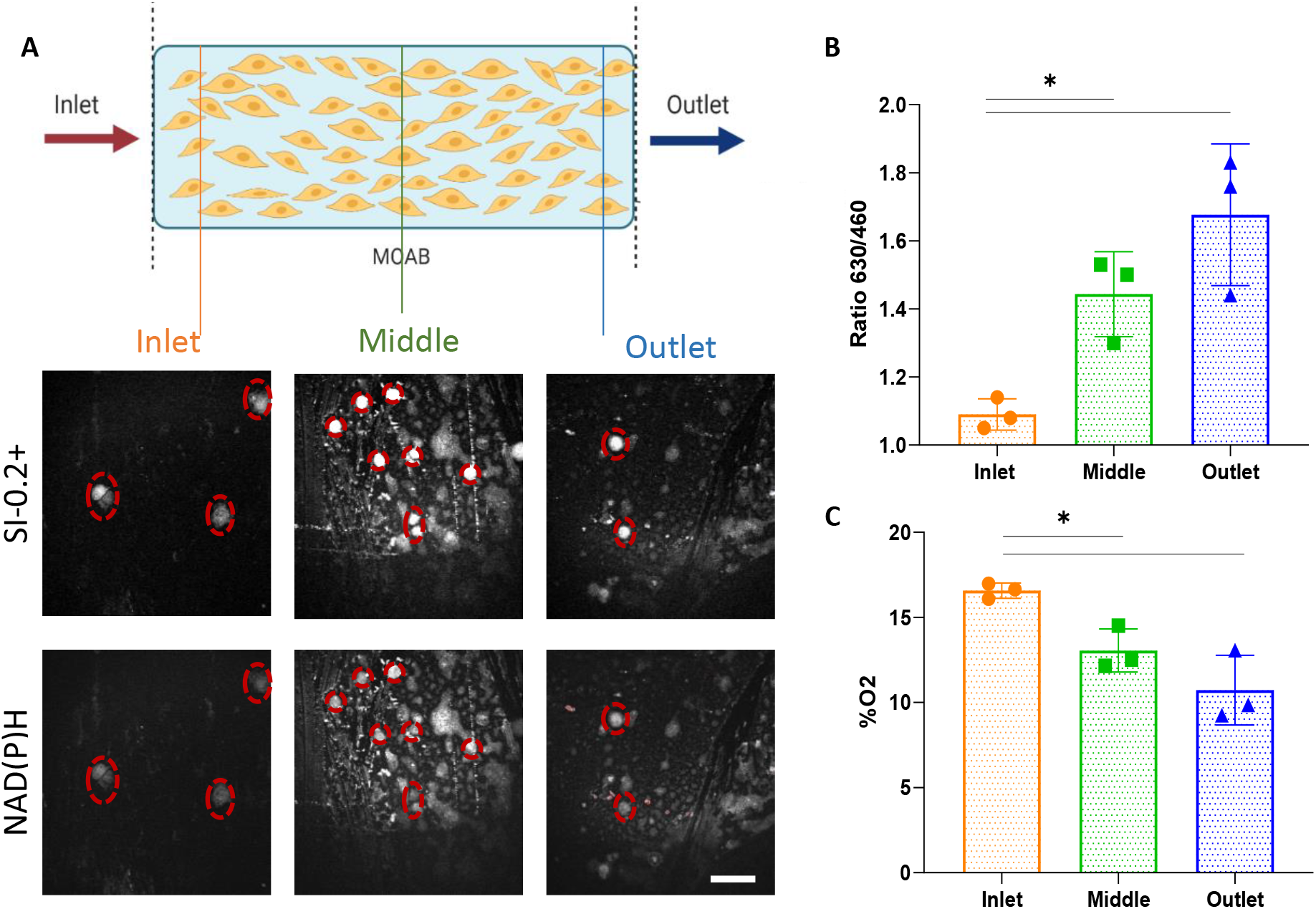
Multiphoton quantitative intracellular oxygen imaging. (A)Representative images acquired at the inlet, middle and outlet areas for both SI-0.2+ and NAD(P)H channels and regions of interest (red) used for quantification (B). Fluorescence intensity ratio of SI-0.2+ per NAD(P)H in the inlet, middle and outlet areas with n=3. Statistical analysis performed using one-way Anova and p<0.05 (C). Intracellular calculated oxygen tension in the different MOAB areas (n=3). One-way ANOVA was used to verify the statistical different with p<0.05.

The effect of downstream decreases of oxygen levels in microfluidic platforms has been previously reported.^58,59^ Mehta et al., using a similar approach of fluorescence intensity measurements have also observed a decreasing trend of oxygen levels from the inlet to outlet areas of a flow bioreactor with a direct dependency on cellular density and the same flow rate of 0.5μl min^−1^ ^58^. Other approaches have employed electrode-based or extracellular fluorescence probes to quantify dissolved oxygen in cell culture medium.^60–63^ However, these approaches lack the opportunity to quantify directly intercellular oxygen concentrations and are more susceptible to experimental error.^58,64^ Moreover, our approach allows direct quantification within microfluidic cultures contained in compact system without the need of complex channels geometries, multilayers structures or inflow of gas mixtures inside the culture chamber.^31^ In order to further assess the energetic state of the cultured h-MSCs, we then employed 2P-FLIM NAD(P)H to directly evaluate the impact this transient oxygen tension on cellular metabolism.

#### Two-photon fluorescence lifetime imaging microscopy (2P-FLIM) evaluation of cellular metabolism

2P-NAD(P)H FLIM was performed on the MOAB on varying sections (similar to the intracellular oxygen measurements): inlet, middle and outlet in order to fully delineate any metabolic heterogeneity of these cells. The MOAB was perfused at the flow rate of 0.5 μL min^−1^ and was imaged at 96 hours of culture growth (Fig. 5A).

**Fig. 5.**
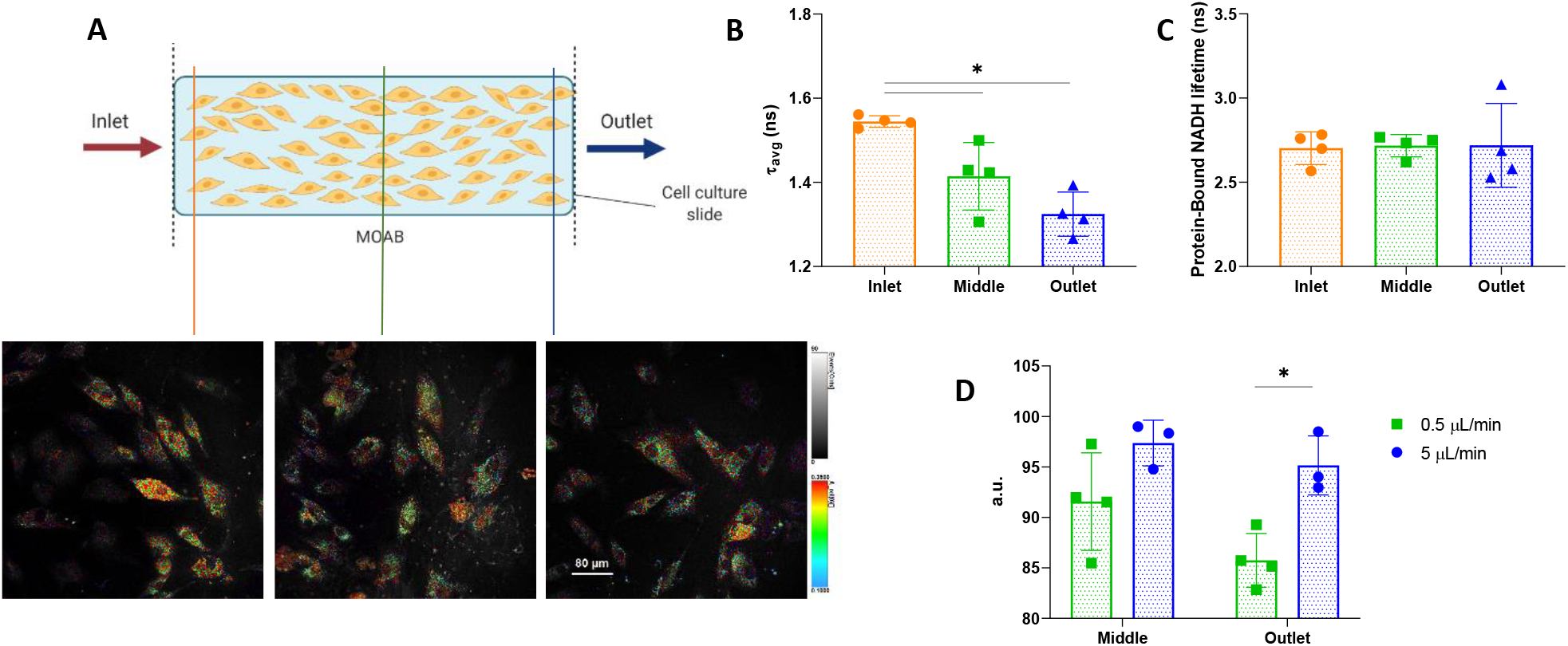
2P-FLIM NAD(P)H imaging of h-MSC cultured on the MOAB. Representative images of the areas imaged after 96 hours cell culture under 0.5 μL min^−1^ flow rate (A). Average fluorescence lifetimes (τ_avg_) calculated for the inlet, middle and outlet areas with n=4. One-way Anova was used to verify statistical difference with p<0.05 (B). Protein-bound NAD(P)H fluorescence lifetime (τ_1_) obtained with n=4 and no statistical difference between groups, verified by One-way Anova (C). Normalisation of middle and outlet τ_avg_ to inlet area of the MOAB at different flow rates. Unpaired t-test was used to verify statistical difference with p<0.05.

FLIM analysis inside of the MOAB in different areas revealed a decreasing gradient in the average fluorescence lifetime while the protein bound NAD(P)H lifetime remains stable. (Fig. 5B,C). The calculated average fluorescence lifetime for the inlet, middle and outlet are 1.545 ± 0.012 ns, 1.415 ± 0.070 ns, 1.325 ± 0.045 ns, respectively. In addition, the middle and outlet values are statistically significantly decreased when compared to inlet values. The calculated average fluorescence lifetime (τ_avg_) has been used regularly in previous works as a unique and easy value to quickly discriminate between cellular metabolic states. ^47,65^ A higher τ_avg_ is reflective of a higher fraction and/or lifetime of the τ_2_ demonstrative of a more OxPhos dependent metabolism. These values decreased as shown on figure 5B, are illustrative of a less dependence on the cell to use OxPhos and therefore becoming more glycolytic.

This trend shows an adaptation of cellular metabolism to the environmental conditions, specifically oxygen; which were predicted to vary in space according to our computational analysis and later measured with our oxygen probe (figure 3,4). As reported on literature, in reduced presence of oxygen the cells adapt from an aerobic to anaerobic metabolism.^6^

The observed trend of decreasing τ_avg_ and its connection to an increase in glycolysis was been reported in several times in literature.^36,47,66^ Walsh et al., demonstrated a decrease in NAD(P)H τ_avg_ in aggressive types of cancer cells, known for being more reliant in glycolysis as an hallmark for their metabolic behaviour.^67,68^ A work by Meleshina et al., showed a contrast between osteogenic differentiated and undifferentiated mesenchymal stem cells. The latter were characterized by a lower NAD(P)H lifetimes associated with their glycolytic metabolic profile.^36^ Therefore, our 2P-FLIM results clearly demonstrate a decrease of τ_avg_ at the outlet of the MOAB chamber which is correlated with higher dependence on the glycolytic pathway by h-MSCs as they are farther away from the fluidic chamber inlet.

However, protein-bound NAD(P)H fluorescence lifetimes (τ_2_) remain stable with 2.703 ± 0.084 ns, 2.718 ± 0.058 ns, 2.719 ± 0.216 ns, for the inlet, middle and outlet, respectively (Figure 5c). According to Blacker et al., τ_2_ is an indicator of the ratio of NADPH and NADH.^69^ These two species are difficult to separate due to spectroscopy properties however, a longer τ_2_ is associated with a higher presence of intracellular NADPH. In the results obtained, these values remain the same which is indicative of a constant ratio of NADPH/NADH pools. NADPH is a metabolic co-factor more involved in anabolic pathways whose presence is less relevant for glycolysis and OxPhos.^50^ Using our 2P-FLIM to distinguish between OxPhos and glycolysis, we can be confident that the OxPhos metabolic pathway is the one more affected by the lack of oxygen, in specific the enzymatic reactions steps that are dependent of oxygen such as the electron transport chain, as already described in literature.^70^

The same experiment was performed with a higher flow rate (5 μL min^−1^) (supplemental figure 2). According to our multiphysics numerical analysis (figure 3), the oxygen tension profile inside the fluidic chamber, in this condition, follows a linear (R^2^ = 0.97) decay, along the principal flow direction, with a higher slope with respect to the 0.5 μL min^−1^ condition. To better understand the changes in cellular metabolism between 5 μL min^−1^ and 0.5 μL min^−1^, a normalisation of the middle and outlet areas to the inlet area was performed. As observed in figure 5C, the average fluorescence lifetimes at 5 μL min^−1^ are higher than the 0.5 μL min^−1^ flowrate being statistically different in the outlet area. With higher values of τ_avg_, the cells experiencing 5 μL min^−1^ flow rate results to be more spatially dependent on OxPhos than in the case of lower flow rate values. In conjugation with the computational mass transport analysis, the non-significant change of τ_avg_ at a higher flow rate can be correlated to the availability of oxygen inside the MOAB. As the cells experience higher oxygen levels in their environment, they will be less reliant on the use of glycolysis.

Our results strongly correlate the increase of glycolytic behavious of with the decrease of oxygen availability predicted *in silico (figure 2)*, which was validated and quantified *in vitro* (figure 3 and 4). The relationship between oxygen availability and metabolism has been widely addressed in many studies where often the metabolic switch promoted by the lack of oxygen is majorly due to the hypoxia-inducible factors (HIFs) ^71–74^. These transcription factors are composed by a stable β-subunit and two α-subunits in which of HIF1α is oxygen-labile.^75,76^ Reducing the levels of oxygen promotes the stabilisation and inactivates the HIF prolyl hydroxylases which results in increased levels of intracellular HIFs.^77,78^ After activation, these factors can promote the expression of genes related with the glycolytic pathway such as: glucose transporter 1 (GLUT1), increasing glucose uptake; lactate dehydrogenase A, converting pyruvate to lactate; and pyruvate dehydrogenase kinase 1 (PDK1) which inhibits the role of pyruvate dehydrogenase reducing the pyruvate uptake by the mitochondria consequently reducing OxPhos and oxygen consumption^79^. Therefore, there is a correlation between the oxygen availability and cellular metabolism, validated by our oxygen measurements and 2P-FLIM.

While there are some reports of FLIM being applied to fluidic systems,^80,81^ this paper demonstrates the first case of 2P-FLIM metabolic characterization of stem cells in a microfluidic bioreactor without the use of microsensors or disruptive end-point assays.^34^ Not only that, this study directly links and validates *in silico* modelling to real-time noninvasive imaging of oxygen concentration gradients followed by its impact on cellular metabolism. The focus of future studies could be the use of 3D substrates with different morphologies in order to understand their mechanical impact on cellular metabolism and the oxygen distribution ^27,82^.

In addition, the real-time, label-free metabolic analysis coupled with the possibility to model oxygen distribution can be used as a more physiological relevant cancer drug screening platform. For example, one major trigger of tumour angiogenesis is a pathological hypoxia achieved due to a decrease in blood flow, reduced vascularization and increase in oxygen consumption ^83^.

## Conclusions

We developed and validated a miniaturized platform for profiling stem cell metabolism in a niche-on-a-chip configuration. Using *in silico* modelling we predicted that this bioreactor could sustain a microphysiological environment with transient spatial oxygen tension distribution typical of the perivascular stem cell niche. This was confirmed by microscopy-based real-time quantification of intracellular oxygenation highlighting a correlating gradient in single cell intracellular oxygen concentration. Finally, we performed high-resolution FLIM measurements of h-MSCs metabolism which allowed us to assess the generation of an oxygen-dependent metabolic profile. A decrease of the average fluorescence lifetime of NAD(P)H is indicative of a higher reliance on glycolysis as the chamber gets more hypoxic towards the outlet and we demonstrated that we can tune the metabolic shift by controlling the environmental oxygen tension through perfusion flowrates and cell density. This toolbox is an innovative *in vitro* platform that can allow tuneable and measurable oxygen tension gradients that can achieve appreciable impacts on cellular metabolism. This advanced transient microenvironment, can recapitulate aspects of intravital niches providing a reliable tool for diseases modelling and drugs screening.

## Materials and Methods

### The microfluidic bioreactor

The MOAB is a miniaturized bioreactor for interstitial flow perfusion and high-resolution imaging. It is fabricated by injection moulding of medical grade PC and consists in one main body and three independent lids that magnetically seal on it. In its closed configuration every lid hosts a rectangular base (3 mm × 6 mm × 0.5 mm, volume = 9 μL) cell culture chamber with a glass surface for cell seeding and real-time imaging. Silicon tubes were perpendicularly connected to the chambers (at the inlet and outlet) allowing for perfusion of cell culture medium. The perfusion was performed by a high precision syringe pump (NE-1600, NEW ERA, Pump Systems Inc.).

The MOAB was maintained at 37°C and 5% CO_2_ during cell culture and fitted on a heating stage to maintain the correct temperature during image acquisition.

### Human Mesenchymal Stem Cells (h-MSCs) culture

Bone marrow derived human-MSCs (h-MSCs) from Lonza^®^ were expanded in Dulbecco’s Modified Eagle’s Medium (DMEM) – Low glucose (Sigma Aldrich), supplemented with 10% Fetal Bovine Serum (FBS) and 1% Penicillin Streptomycin (Pen/Strep). h-MSCs at P3 were seeded in each of the three MOAB’s culture chambers, after overnight pre-conditioning in complete DMEM, at a cell density of 3.5 × 10^4^ cells/cm^2^. The culture chambers were incubated at 37°C and 5% CO_2_ for 3 hours to allow h-MSCs adhering to the surface before beginning flow perfusion.

### Mathematical modelling

A Computational Fluid Dynamic (CFD) analysis was performed to investigate the velocity profile and shear stress distribution inside the cell culture chamber at the cellular level.

A 3D, stationary model was created in COMSOL Multiphysics^®^ 5.2 (Burlington, MA, USA). The medium flow through the micro bioreactor was simulated by implementing the Navier-Stokes equation and the continuity equation for local conservation of momentum and mass, respectively. The flow was assumed laminar and fully developed at the entrance of the chamber with inlet velocity values computed starting from the selected flowrate range (0.5-5 μL min^−1^). The mass transfer model assumed MM rate of uptake of oxygen by the cells.

**Table 1.** Model parameters and values used for the computational analysis

|  |  |  |  |  |
| --- | --- | --- | --- | --- |
| Oxygen diffusion coefficient | DO<sub>2</sub> | 2x10<sup>-9</sup> | m<sup>2</sup> s<sup>-1</sup> | 84 |
| Dynamic viscosity | η | 8.4x10<sup>-4</sup> | Pa s | 84 |
| Michaelis-Menten constant for oxygen uptake | K<sub>m,O</sub> | 4.05x10<sup>-9</sup> | mol cm<sup>-3</sup> | 52 |
| Maximum uptake rate of oxygen | V<sub>max</sub> | 100 | fmol h<sup>-1</sup> cell<sup>-1</sup> | 52 |

### Quantitative intracellular O_2_ imaging

We measured intracellular O_2_ tension by using conjugated polymer nanoparticles for high-resolution O_2_ imaging in living cells. The nanoparticle probes (bead name: Si-02^+^) are constituted by a substituted conjugated polymer (polyfluorene) covalently bounded with a phosphorescent metalloporphyrin which emits in far red wavelengths, in a low oxygen environment, due to Fluorescence Resonance Energy Transfer (FRET) mechanism. Details on the manufacturing and fluorescence properties of the copolymer complexes can be found in ^57^. After seeding h-MSCs inside the MOAB, the staining with nanoparticles was achieved by incubation at 8 μg/mL for 16 hours in order to allow for probes internalization. The culture chambers were washed twice with DMEM medium. Imaging and related measurements were performed, after 96 hours perfusion at 0.5 μL min^−1^, with two-photon microscope (Olympus BX61WI) in phenol red-free DMEM.

The excitation wavelength was set at 760 nm and emission wavelength signal was detected in two bandpass filters at 455-490 nm for the fluorene backbone (460 nm) and 580-638 nm for the PT(II)-Porphyrin complex (630 nm) ^57^.

In order to correlate the fluorescence intensity signal with the intracellular oxygen tension, a semi-calibration curve was obtained (supplemental figure 1). To achieve this, stained h-MSCs were imaged inside of the MOAB in static condition to ensure an oxygen homogenisation. First, the stained h-MSCs were imaged in normoxia environment followed, in a separate experiment, by imaging in hypoxia conditions with the respiratory ability of the cells inhibited. To achieve this, cells were incubated at 37C for 1 hour using 2.5 μM of Rotenone and 10 μM Antimycin A as these metabolic inhibitors act upon complex I and III of the electron transport chain, negating the cell ability to consume oxygen. Using the same excitation power, and photomultiplier (PMT) detector settings, h-MSCs cells grown in a MOAB device for 3 days at 0.5ul/min flow rate. Afterwards, they were stained and imaged using the multiphoton microscope. Imaging analysis was performed using ImageJ to select ROI areas and calculate their mean gray value.

### 2-Photon microscopy and fluorescence lifetime imaging analysis

2P-FLIM was performed using a custom upright (Olympus BX61WI) laser multiphoton microscopy system equipped with Titanium: sapphire laser (Chameleon Ultra, Coherent^®^, USA), water-immersion 25x objective (Olympus, 1.05NA) and temperature controlled stage at 37°C. During the measurements, the syringe pump was working with the flow-rate adequate to the experiment being performed (5ul/min or 0.5ul/min) or in static conditions for the O_2_ probe calibration measurements.

Two photon excitation of NAD(P)H fluorescence was performed at the excitation wavelength of 760 nm. A 455/90 nm bandpass filter was used to isolate the NAD(P)H fluorescence emission. 512 × 512 pixel images were acquired with a pixel dwell time of 3.81 μs and 30 s collection time. A PicoHarp 300 TCSPC system operating in the time-tagged mode coupled with a photomultiplier detector assembly (PMA) hybrid detector (PicoQuanT GmbH, Germany) was used for fluorescence decay measurements yielding 256 time bins per pixel.

Fluorescence lifetime images with their associated decay curves for NAD(P)H were obtained and region of interest (ROI) analysis of the total cells present on the image was performed in order to remove any background artefact. The decay curved was generated and was fitted with a double-exponential decay without including the instrument response function (IRF) (Eq.1).

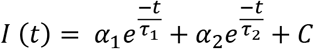

I(t) corresponds to the fluorescence intensity measured at time t after laser excitation; α_1_ and α_2_ represents the fraction of the overall signal proportion of a short and long component lifetime components, respectively. τ_1_ and τ_2_ are the short and long lifetime components, respectively; C corresponds to background light. χ2 statistical test was used to evaluate the goodness of multi-exponential fit to the raw fluorescence decay data. In this study all the values with χ2 < 1.3 were considered as ‘good’ fits.

For NAD(P)H, the double exponential decay was used to differentiate between the free (τ_1_) and protein-bound (τ_2_) NAD(P)H. The average fluorescence lifetime was calculated using equation 2.

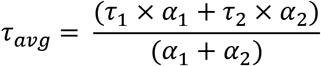

### Statistical analysis

The data obtained in the course of several experiments was evaluated for statistical significance as described in the figure descriptions. Paired t-test or One-way ANOVA (where appropriate) were used to establish statistical differences when p-value was lower than 0.05.

## Author contributions

MGM, MTR, NN, SP and CdN designed experiments and wrote the paper. NN, SP and CdN performed experiments, collected and analysed the data. MGM and MTR revised the manuscript.

## Acknowledgements

The author would like to acknowledge the contribution of Dr. Ruslan Dmitriev for providing guidance and the fluorescence probes for the intracellular oxygen measurements performed.

This research has received funding from: the European Research Council (ERC) under the European Union’s Horizon 2020 research and innovation program (G.A. No. 825159 – MOAB); the Italian Ministry of University and Research (MUR) under the grant program MIUR-FARE-2016 (G.A. No. R16ZNN2R9K – BEYOND); the National Centre for the Replacement, Refinement and Reduction of Animals in Research (NC3Rs) under the CleanCut 2020 Challenge (G.A. No. NC/C01903/1 – MOAB). NN is supported by a Trinity College Dublin, Provost’s PhD Award and the TCD FLIM core unit directed by MM is supported by a SFI Infrastructure Programme: Category D Opportunistic Funds Call

## References

1 M. Crisan, M. Corselli, W. C. W. Chen and B. Péault, J. Cell. Mol. Med., 2012, 16, 2851–2860.

2 M. Oh and J. E. Nör, Front. Physiol., 2015, 6.

3 D. E. Discher, D. J. Mooney and P. W. Zandstra, Science (80-.)., 2009, 324, 1673–1677.

4 S. Méndez-Ferrer, D. Bonnet, D. P. Steensma, R. P. Hasserjian, I. M. Ghobrial, J. G. Gribben, M. Andreeff and D. S. Krause, Nat. Rev. Cancer, 2020, 20, 285–298.

5 A. Benabid and L. Peduto, Curr. Opin. Immunol., 2020, 64, 50–55.

6 M. G. Vander Heiden, L. C. Cantley and C. B. Thompson, Science (80-.)., 2009, 324, 1029–1033.

7 N. Shyh-Chang, G. Q. Daley and L. C. Cantley, Dev., 2013, 140, 2535–2547.

8 S. Palomäki, M. Pietilä, S. Laitinen, J. Pesälä, R. Sormunen, P. Lehenkari and P. Koivunen, Stem Cells, 2013, 31, 1902–1909.

9 C. Hu, L. Fan, P. Cen, E. Chen, Z. Jiang and L. Li, Int. J. Mol. Sci., 2016, 17.

10 K. Ito and K. Ito, Annu. Rev. Cell Dev. Biol., 2016, 32, 399–409.

11 L. da Silva Meirelles, P. C. Chagastelles and N. B. Nardi, J. Cell Sci., 2006, 119, 2204–2213.

12 Y. Kfoury and D. T. Scadden, Cell Stem Cell, 2015, 16, 239–253.

13 B. Fattizzo, J. A. Giannotta and W. Barcellini, Int. J. Mol. Sci., 2020, 21.

14 C. M. Rivera-Cruz, J. J. Shearer, M. Figueiredo Neto and M. L. Figueiredo, Stem Cells Int., 2017, 2017.

15 M. Tewary, N. Shakiba and P. W. Zandstra, Nat. Rev. Genet., 2018, 19, 595–614.

16 X. Yuan, T. M. Logan and T. Ma, Front. Immunol., 2019, 10, 977.

17 K. Bai, Y. Huang, X. Jia, Y. Fan and W. Wang, J. Biomech., 2010, 43, 1176–1181.

18 B. Trappmann, J. E. Gautrot, J. T. Connelly, D. G. T. Strange, Y. Li, M. L. Oyen, M. A. C. Stuart, H. Boehm, B. Li, V. Vogel, J. P. Spatz, F. M. Watt and W. T. S. Huck, Nat. Mater., 2012, 11, 1–8.

19 J. K. Leach and J. Whitehead, ACS Biomater. Sci. Eng., 2017, acsbiomaterials.6b00741.

20 K. M. Kim, Y. J. Choi, J.-H. Hwang, A. R. Kim, H. J. Cho, E. S. Hwang, J. Y. Park, S.-H. Lee and J.-H. Hong, PLoS One, 2014, 9, e92427.

21 D. Mushahary, A. Spittler, C. Kasper, V. Weber and V. Charwat, Cytom. Part A, 2018, 93, 19–31.

22 T. J. Bartosh and J. H. Ylostalo, Cells,, DOI:10.3390/cells8091031.

23 K. Carter, H. J. Lee, K. S. Na, G. M. Fernandes-Cunha, I. J. Blanco, A. Djalilian and D. Myung, Acta Biomater., 2019, 99, 247–257.

24 Y. S. Torisawa, C. S. Spina, T. Mammoto, A. Mammoto, J. C. Weaver, T. Tat, J. J. Collins and D. E. Ingber, Nat. Methods, 2014, 11, 663–669.

25 S. Sieber, L. Wirth, N. Cavak, M. Koenigsmark, U. Marx, R. Lauster and M. Rosowski, J. Tissue Eng. Regen. Med., 2018, 12, 479–489.

26 C. Lin, L. Lin, S. Mao, L. Yang, L. Yi, X. Lin, J. Wang, Z. X. Lin and J. M. Lin, Anal. Chem., 2018, 90, 10326–10333.

27 A. Marturano-Kruik, M. M. Nava, K. Yeager, A. Chramiec, L. Hao, S. Robinson, E. Guo, M. T. Raimondi and G. Vunjak-Novakovic, Proc. Natl. Acad. Sci. U. S. A., 2018, 115, 1256–1261.

28 Y. S. Torisawa, C. S. Spina, T. Mammoto, A. Mammoto, J. C. Weaver, T. Tat, J. J. Collins and D. E. Ingber, Nat. Methods, 2014, 11, 663–669.

29 M. D. Brennan, M. L. Rexius-Hall, L. J. Elgass and D. T. Eddington, Lab Chip, 2014, 14, 4305–4318.

30 A. Super, N. Jaccard, M. P. Cardoso Marques, R. J. Macown, L. D. Griffin, F. S. Veraitch and N. Szita, Biotechnol. J., 2016, 11, 1179–1189.

31 K. R. Rivera, M. A. Yokus, P. D. Erb, V. A. Pozdin and M. Daniele, Analyst, 2019, 144, 3190–3215.

32 A. Carreau, B. El Hafny-Rahbi, A. Matejuk, C. Grillon and C. Kieda, J. Cell. Mol. Med., 2011, 15, 1239–1253.

33 A. Mohyeldin, T. Garzón-Muvdi and A. Quiñones-Hinojosa, Cell Stem Cell, 2010, 7, 150–161.

34 J. Kieninger, A. Weltin, H. Flamm and G. A. Urban, Lab Chip, 2018, 18, 1274.

35 K. Suhling, L. M. Hirvonen, J. A. Levitt, P. Chung, C. Tregidgo, A. Le, D. A. Rusakov, K. Zheng, S. Ameer-beg, S. Poland, S. Coelho, R. Henderson and N. Krstajic, Chem. Phys. Lett., 2015, 27, 3–40.

36 A. V Meleshina, V. V Dudenkova, A. S. Bystrova, D. S. Kuznetsova, M. V Shirmanova and E. V Zagaynova, Stem Cell Res. Ther., 2017, 1–10.

37 L. Izzo, M. Tunesi, L. Boeri, M. Laganà, C. Giordano and M. T. Raimondi,, DOI:10.1007/s10544-019-0387-8.

38 A. B. Yeatts, D. T. Choquette and J. P. Fisher, Biochim. Biophys. Acta - Gen. Subj., 2013, 1830, 2470–2480.

39 E. Stavenschi, M. Labour and D. A. Hoey, J. Biomech., 2017, 55, 99–106.

40 M. Tonin, N. Descharmes and R. Houdré, 2014, 16, 465.

41 T. S. Blacker and M. R. Duchen, Free Radic. Biol. Med., 2016, 100, 53–65.

42 J. M. Szulczewski, D. R. Inman, D. Entenberg, S. M. Ponik, J. Aguirre-Ghiso, J. Castracane, J. Condeelis, K. W. Eliceiri and P. J. Keely, Sci. Rep., 2016, 6, 25086.

43 J. R. Lakowicz, H. Szmacinski, K. Nowaczyk and M. L. Johnson, Proc. Natl. Acad. Sci., 1992, 89, 1271–1275.

44 K. P. Quinn, E. Bellas, N. Fourligas, K. Lee, D. L. Kaplan and I. Georgakoudi, Biomaterials, 2012, 33, 5341–5348.

45 A. Varone, J. Xylas, K. P. Quinn, D. Pouli, G. Sridharan, M. E. McLaughlin-Drubin, C. Alonzo, K. Lee, K. Munger and I. Georgakoudi, Cancer Res, 2014, 74, 3067–3075.

46 P. H. Lakner, M. G. Monaghan, Y. Möller, M. A. Olayioye and K. Schenke-Layland, Sci. Rep., 2017, 7, 42730.

47 I. A. Okkelman, N. Neto, D. B. Papkovsky, M. G. Monaghan and R. I. Dmitriev, Redox Biol, 2019, 30, 101420.

48 M. C. Skala, K. M. Riching, D. K. Bird, A. Gendron-Fitzpatrick, J. Eickhoff, K. W. Eliceiri, P. J. Keely and N. Ramanujam, J. Biomed. Opt., 1992, 12, 024014.

49 M. C. Skala, K. M. Riching, A. Gendron-Fitzpatrick, J. Eickhoff, K. W. Eliceiri, J. G. White and N. Ramanujam, Proc Natl Acad Sci U S A, 2007, 104, 19494–19499.

50 W. Ying, Antioxid Redox Signal, 2008, 10, 179–206.

51 J. M. Rutkowski and M. A. Swartz, Trends Cell Biol., 2007, 17, 44–50.

52 M. J. Osiecki, S. D. L. McElwain and W. B. Lott, PLoS One,, DOI:10.1371/journal.pone.0202079.

53 M. T. Raimondi, C. Giordano and R. Pietrabissa, J. Appl. Biomater. Funct. Mater., 2015, 13, e313–e319.

54 M. Crisan, S. Yap, L. Casteilla, C. W. Chen, M. Corselli, T. S. Park, G. Andriolo, B. Sun, B. Zheng, L. Zhang, C. Norotte, P. N. Teng, J. Traas, R. Schugar, B. M. Deasy, S. Badylak, H. J. Buhring, J. P. Giacobino, L. Lazzari, J. Huard and B. Péault, Cell Stem Cell, 2008, 3, 301–313.

55 I. Özen, J. Boix and G. Paul, Perivascular mesenchymal stem cells in the adult human brain: a future target for neuroregeneration?, 2012, vol. 1.

56 D. B. Papkovsky, · Ruslan and I. Dmitriev, 2018, 75, 2963–2980.

57 R. I. Dmitriev, S. M. Borisov, H. Dü, S. Sun, B. J. Mü, J. Prehn, V. P. Baklaushev, I. Klimant and D. B. Papkovsky,, DOI:10.1021/acsnano.5b00771.

58 G. Mehta, K. Mehta, D. Sud, J. W. Song, T. Bersano-Begey, N. Futai, Y. Seok Heo, M.-A. Mycek, J. J. Linderman, S. Takayama, G. Mehta, · D Sud, J. W. Song, · T Bersano-Begey, · N Futai, Y. S. Heo, M.-A. Mycek, J. J. Linderman, S. Takayama, K. Mehta and · J J Linderman, Biomed Microdevices, 2007, 9, 123–134.

59 J. W. Allen, Toxicol. Sci., 2005, 84, 110–119.

60 S. Andreescu, O. A. Sadik, D. W. McGee and S. Suye, Anal. Chem., 2004, 76, 2321–2330.

61 T. H. Park and M. L. Shuler, Biotechnol. Prog., 2003, 19, 243–253.

62 J. Malda, J. Rouwkema, D. E. Martens, E. P. le Comte, F. K. Kooy, J. Tramper, C. A. van Blitterswijk and J. Riesle, Biotechnol. Bioeng., 2004, 86, 9–18.

63 W. Zhong, P. Urayama and M.-A. Mycek, J. Phys. D. Appl. Phys., 2003, 36, 1689–1695.

64 I. R. Sweet, G. Khalil, A. R. Wallen, M. Steedman, K. A. Schenkman, J. A. Reems, S. E. Kahn and J. B. Callis, Diabetes Technol. Ther., 2002, 4, 661–672.

65 A. J. Walsh, K. P. Mueller, K. Tweed, I. Jones, C. M. Walsh, N. J. Piscopo, N. M. Niemi, D. J. Pagliarini, K. Saha and M. C. Skala, Nat. Biomed. Eng.,, DOI:10.1038/s41551-020-0592-z.

66 A. J. Walsh, R. S. Cook, H. C. Manning, D. J. Hicks, A. Lafontant, C. L. Arteaga and M. C. Skala, Cancer Res, 2013, 73, 6164–6174.

67 A. Walsh, R. S. Cook, B. Rexer, C. L. Arteaga and M. C. Skala, Biomed Opt Express, 2012, 3, 75–85.

68 M. G. Vander Heiden and R. J. DeBerardinis, Cell, 2017, 168, 657–669.

69 T. S. Blacker, Z. F. Mann, J. E. Gale, M. Ziegler, A. J. Bain, G. Szabadkai and M. R. Duchen, Nat. Commun.,, DOI:10.1038/ncomms4936.

70 Y. Liu, G. Fiskum and D. Schubert, J. Neurochem., 2002, 80, 780–787.

71 L. A. J. O’Neill, R. J. Kishton and J. Rathmell, Nat. Rev. Immunol., 2016, 16, 553–565.

72 S. Y. Lunt and M. G. Vander Heiden, Annu. Rev. Cell Dev. Biol., 2011, 27, 441–464.

73 K. L. Eales, K. E. R. Hollinshead and D. A. Tennant, Oncogenesis, 2016, 5, e190–e190.

74 R. D. Guzy and P. T. Schumacker, Exp. Physiol., 2006, 91, 807–819.

75 W. Luo, H. Hu, R. Chang, J. Zhong, M. Knabel, R. O’Meally, R. N. Cole, A. Pandey and G. L. Semenza, Cell, 2011, 145, 732–744.

76 G. L. Wang, B. H. Jiang, E. A. Rue and G. L. Semenza, Proc. Natl. Acad. Sci., 1995, 92, 5510–5514.

77 A. C. R. Epstein, J. M. Gleadle, L. A. McNeill, K. S. Hewitson, J. O’Rourke, D. R. Mole, M. Mukherji, E. Metzen, M. I. Wilson, A. Dhanda, Y.-M. Tian, N. Masson, D. L. Hamilton, P. Jaakkola, R. Barstead, J. Hodgkin, P. H. Maxwell, C. W. Pugh, C. J. Schofield and P. J. Ratcliffe, Cell, 2001, 107, 43–54.

78 B. H. Jiang, G. L. Semenza, C. Bauer and H. H. Marti, Am. J. Physiol. Physiol., 1996, 271, C1172–C1180.

79 W. W. Wheaton and N. S. Chandel, Am. J. Physiol. Physiol., 2011, 300, C385–C393.

80 N. Shen, J. A. Riedl, D. A. Carvajal Berrio, Z. Davis, M. G. Monaghan, S. L. Layland, S. Hinderer and K. Schenke-Layland, Biomed. Mater., 2018, 13, 24101.

81 H. M. Wu, T. A. Lee, P. L. Ko, W. H. Liao, T. H. Hsieh and Y. C. Tung, Analyst, 2019, 144, 3494–3504.

82 A. Remuzzi, B. Bonandrini, M. Tironi, L. Longaretti, M. Figliuzzi, S. Conti, T. Zandrini, R. Osellame, G. Cerullo and M. T. Raimondi, Cells,, DOI:10.3390/cells9081873.

83 C. T. Taylor and S. P. Colgan, Nat. Rev. Immunol., 2017, 17, 774–785.

84 M. Laganà and M. T. Raimondi, 2012, 225–234.

