## Supplemental for "Intracellular label-free detection of mesenchymal stem cell metabolism within a perivascular niche-on-a-chip"

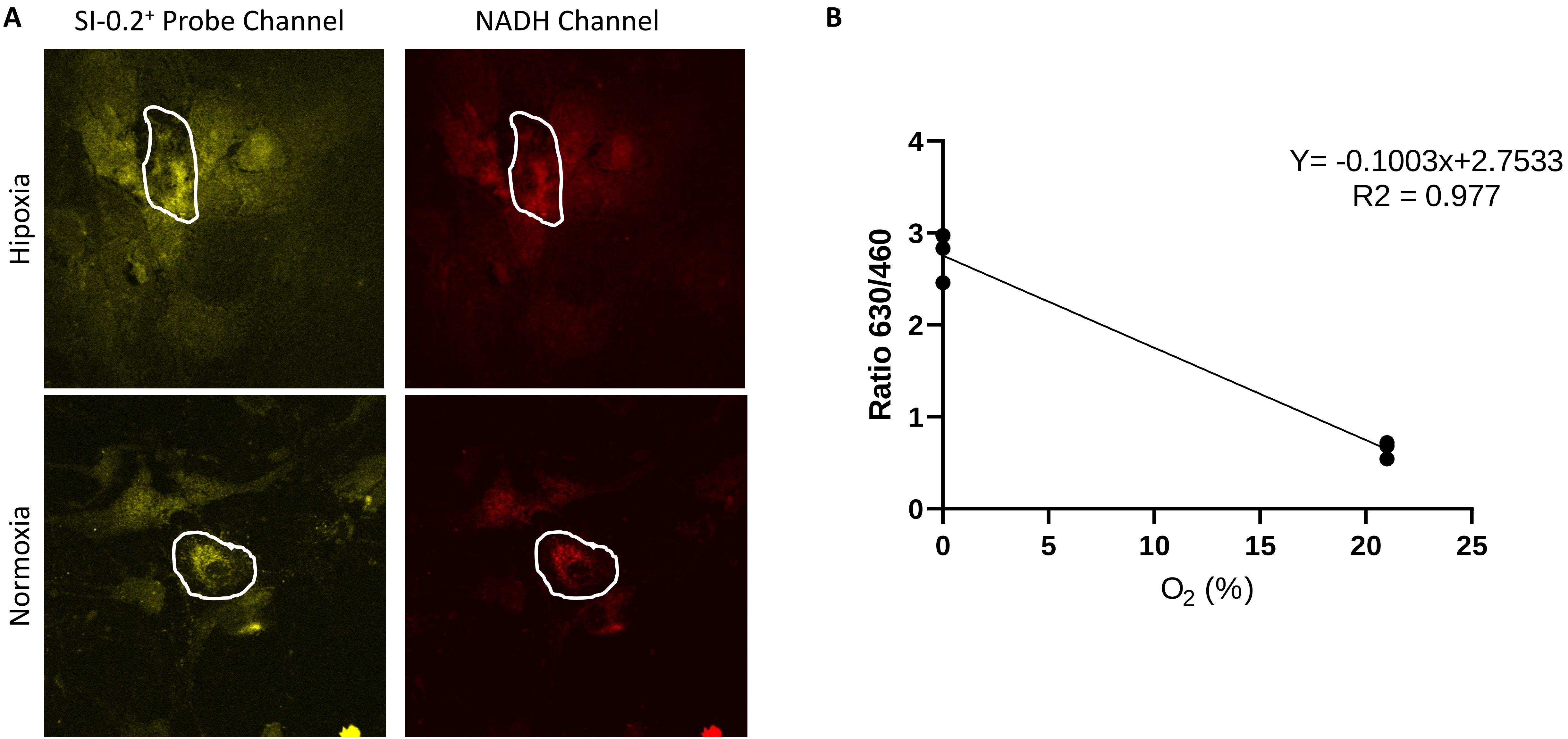


Supplemental Figure 1 – Fluorescence intensity measurements of intracellular oxygen. A) Representative images of hMSC acquired in Normoxia (21%) and Hipoxia (0%) conditions. B) Linear regression and coefficient of determination calculated for the 630/460 ratio measured from ROI areas (n=3). At least 3 images for each n-value were obtained.

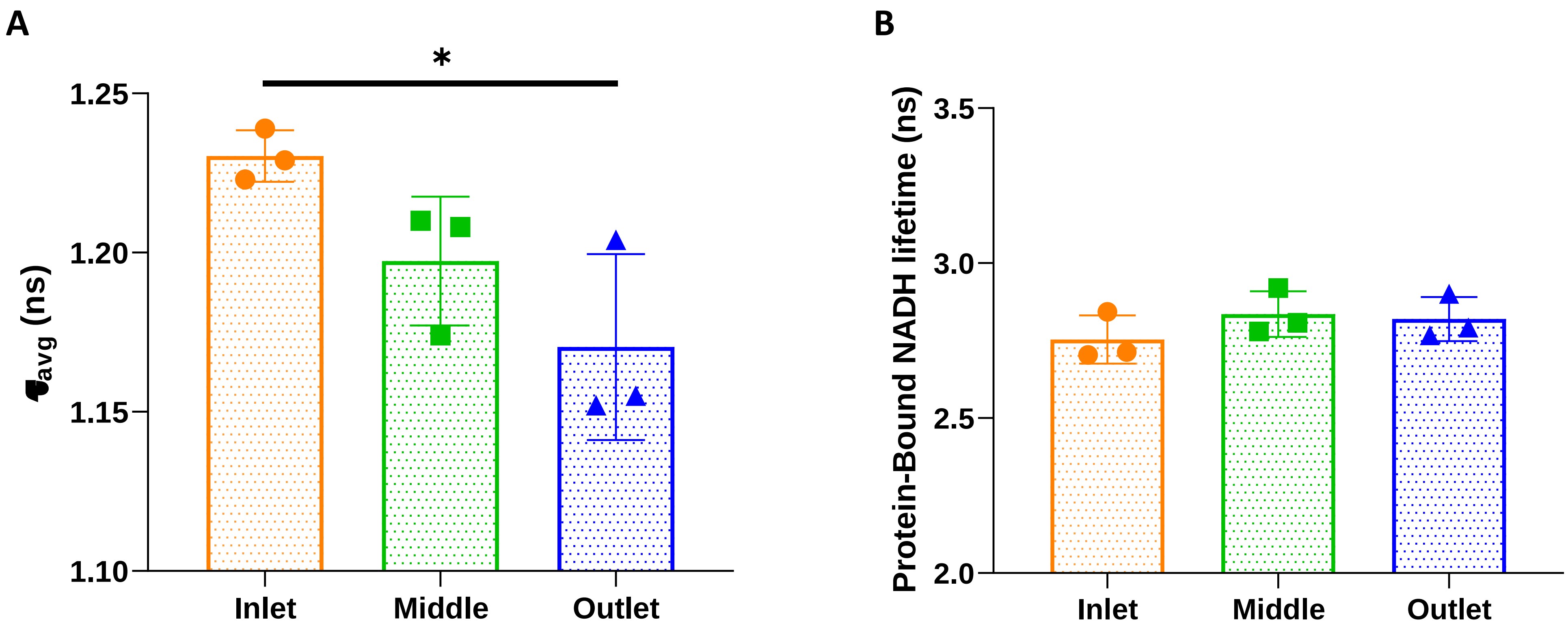


Supplemental Figure 2 – 2P-Photon FLIM of NAD(P)H of hMSC cultured in the MOAB with 5µL $min^{-1}$ . A) Calculated average fluorescence lifetimes for the inlet, middle and outlet region. B) Protein-bound lifetimes measured for the inlet, middle and outlet region. n=3. One-Way Anova was used for to verify statistical difference with p-value < 0.05
